# Neuron synchronization analyzed through spatial-temporal attention

**DOI:** 10.1101/2024.07.10.602834

**Authors:** Haoming Yang, KC Pramod, Panyu Chen, Hong Lei, Simon Sponberg, Vahid Tarokh, Jeffrey A. Riffell

**Affiliations:** Department of Electrical and Computer Engineering, Duke University; Department of Biology, University of Washington, Seattle; Department of Computer Science, Duke University; School of Life Sciences, Arizona State University; Schools of Physics and Biological Sciences, Georgia Institute of Technology

## Abstract

Across diverse organisms, the temporal dynamics of spiking responses between neurons, the neural synchrony, is crucial for encoding different stimuli. Neural synchrony is especially important in the insect antennal (olfactory) lobe (AL). Previous studies on synchronization, however, rely on pair-wise synchronization metrics including the cross-correlogram and cos-similarity between kernelized spikes train. These pair-wise analyses overlook an important aspect of synchronization which is the interaction at the population neuron level. There are also limited modeling techniques that incorporate the synchronization between neurons in modeling population spike trains. Inspired by recent advancements in machine learning, we leverage a modern attention mechanism to learn a generative normalizing flow that captures neuron population synchronization. Our method not only reveals the spiking mechanism of neurons in the AL region but also produces semi-interpretable attention weights that characterize neuron interactions over time. These automatically learned attention weights allow us to elucidate the known principles of neuron synchronization and further shed light on the functional roles of different cell types (the local interneurons (LNs), and projection neurons (PNs)) in the dynamic neural network in the AL. By varying the balance of excitation and inhibition in this neural circuit, our method further uncovers the pattern between the strength of synchronization and the ratio of an odorant in the mixture.

**Author Summary:** The olfactory system can accurately compute the mixture of volatile compounds emitted from distant sources, enabling the foraging species to exhibit fast and effective decisions. However, altering ratios of one of the compounds in the mixture could be perceived as a different odor. Leveraging the current understanding of neural synchronization on sensory neural regions of insects, we construct a spatial-temporal attention normalizing flow, which partially replicates the AL region’s functionality by learning the spiking mechanics of neurons. Beyond providing insights of the spiking mechanism of neurons in the AL region, our method also produces semi-interpretable attention weights that characterize neuron interaction over time. These automatically learned attention weights allow us to dissect out the principles of neuron synchronization and interaction mechanisms between projection neurons (PNs) and local neurons (LNs). Utilizing our accurate model of these AL functionality, we show evidence that the behavioral relevant compounds are closely clustered together while varying the intensities of one of the behavioral compounds in the mixture could attenuate the synchronization

## 1 Introduction

The brain constructs a meaningful perception of the sensory features of the complex external world, such as intensity, modality-specific classification, and stimulus dynamics [1, 2]. This is particularly true in olfaction, where the complex milieu of volatile odor chemicals constantly varies over many orders of magnitude in concentration in the environment, and the scent from a single odor source can be composed of tens to hundreds of compounds. In vertebrate and invertebrate animals, the olfactory system can accurately compute the mixture of volatile compounds emitted from distant sources, enabling the foraging species to exhibit fast and effective decisions [3–5]. This process incorporates different populations of neurons in succession to encode olfactory information into the spiking language of the neural activity such that the relevant odor information is precisely extracted from the mixtures [6].

Olfactory information from the external world is relayed by the spatiotemporal activity of different populations of neurons in the primary olfactory center, the antennal lobe (AL) in insects, and the olfactory bulb (OB) in vertebrates, leading to the stereotypic activation of glomeruli and thus facilitating an important role in odor classification and encoding [7, 8]. The AL in Lepidopteran moths consists of the network of an excitatory afferent axons of olfactory sensory neurons (OSNs, 330,000 in number, [9, 10]), projection neurons (PNs, around 1100 in number, [11]) that relay information from the AL to the higher brain areas, and inhibitory local neurons (LNs, approximately 360, [12]) that form a dense network of interconnecting glomeruli within the AL. Previous studies have shown that the PNs, originating from a single glomerulus, can be excited by the stimulation with one or a few components in the mixture, and inhibited by or unresponsive to the other components. The combination of excitatory and inhibitory components in the mixture, and their relative proportions, may modulate the neural activity [13–16]. The responses of the PNs is mediated by the balance of excitatory drive from the OSNs and inhibition from the activity of the LNs [17, 18].

Previous work has suggested that micro-circuits in the AL, and particularly the temporal coordination of LN and PN spiking responses, including neural spiking synchrony, provide odor-specific representations and encoding. Although neural synchrony between PN pairs has been shown to be modulated by the proportions of components in binary mixtures, how the neural population represents the composition of complex odors, and proportion of compounds in those odors, has been demonstrated only rarely. There is a current need for computational tools to model the spatiotemporal dynamics of the neural ensemble, and characterize how different cell types within the population affect the olfactory representations.

The Sphinx moth *Manduca sexta* is an excellent model for understanding how complex odor information is encoded in the olfactory system. Using its sense of smell, this moth navigates over kilometers to locate patches of hostplant, the *Datura wrightii* flowers. Rather than encoding all 60-80 compounds in the bouquet, it detects around nine critical odorants in the mixture emitted from the flower that elicits its ability to navigate and locate the odor source [3, 5]. Single odorants rarely have behavioral significance to foraging insects. However, a small subset of odorants in the mixture is critical in the odor encoding [3]. Previous studies have shown that changing the concentration of one of the critical compounds significantly affects the moth’s ability to discriminate and track the floral odor [3]. However, the cellular and computational bases by which the olfactory system binds specific features of the complex odor mixture - including the critical odorants - are not known, and neuromorphic principles that are involved in such processes are still uncertain, given the diverse physiological and morphological properties of these neuron types [19–21]. Evaluating the dynamic changes in the AL network in reaction to different concentration effects remains difficult through direct physiological experiments as the quantity delivered to test species is not under the direct control of experiment; therefore, we propose to analyze the behavior of this elaborated AL network through a data-driven, machine-learning perspective.

Previous studies on spatiotemporal encoding of AL neurons have often focused on the spiking synchrony between pairs of neurons [3, 14]. However, biologically, the spatiotemporal responses of the neural population will activate and interact through time. Due to methodological constraints, these previous studies did not capture this important population-level activity. Plasticity in the spiking activities of the neurons in AL is also hard to capture putatively through sensory experiences and inter-subject variabilities. The challenges in modeling the spatiotemporal dynamics at the population level motivated us to build a modeling system to learn the spiking process of a neuron through its history and the related activity of other neurons. The modeling of the neuron spiking process is not new. Generalized linear models with Poisson link function have been well studied for this type of modeling [22]. Latent dynamic methods have also been widely applied to model the spiking activities [23, 24]. However, these methods assume the spike trains follow a temporal process with known arrival time distributions. Recent advancements in deep learning methods that model temporal point processes allow the learning of these processes without assuming a canonical distribution for events’ arrival time. For example, excursions of an Itô’s process have been exploited to learn temporal point process [25].

In this work, we treat the spiking of neurons as a temporal point process. Instead of assuming the arrival time of spikes follows a canonical distribution (e.g. Poisson distribution), we use the highly flexible, non-parametric, deep normalizing flow to model the probability distribution of inter-spike intervals (ISIs) [26]. During the modeling process of the spike train of a specific neuron in the AL, we introduce a novel spatial-temporal attention module to learn how individual neurons synchronize with the rest of the neuron population (spatial) and are affected by population spike trains dynamically (temporal). This spatial attention weight module accounts for the higher order interactions across a population of neurons, allowing us to analyze complex population-level synchronization beyond the pairwise analyses of Ensemble Synchronization Index [3] and Kernelized binless methods [14].

The attention module also allows us to investigate the interaction between different cell types in the microcircuit, including the PNs and LNs. We show that taking all the PNs in an ensemble depicts the discrimination of the behavioral and non-behavioral odor stimuli whereas the LNs did not show significant separation of odor, suggesting that LN mainly modulate the synchronization of PNs. Lastly, we explore how changing the excitatory-inhibitory balance - by changing the proportion of a critical odorant in the natural behavioral stimulus - alters the synchronized activity and odor encoding. The result indicates that increasing the proportion of a compound in the mixture could attenuate the pattern of neural synchrony.

## 2 Data Curation

### 2.1 Insect preparation

Adult male moths (*Manduca sexta*; Lepidoptera: Sphingidae) were reared in the laboratory on an artificial diet under a long-day (17/7-h light/dark cycle) photoperiod. The moths (3 days old, post eclosion) were secured in a 10 ml plastic pipette (Thermofisher Scientific, USA) with dental wax (Kerr Corporation, Romulus, MI, USA) leaving the head and antennae exposed. The cuticle on the head was carefully cut to expose the brain, and all the muscles, trachea, and neural sheath were carefully removed with fine forceps (Fine Science tool, USA). The restrained moth was mounted to a recording platform attached to the vibration–isolation table. The preparation was placed such that the ALs are orientated dorsofrontally. The brain was superfused slowly with physiological saline solution [150 mM NaCl, 3 mM CaCl2, 3 mM KCl, 10 mM N-Tris(hydroxymethyl) methyl-2 aminoethanesulfonic acid buffer, and 25 mM sucrose, pH 6.9] throughout the experiment.

### 2.2 Odor Stimulation

Pulses of air (100ml/min) were pushed through a glass cartridge containing a piece of Whatman filter paper (Millipore sigma, USA) loaded with 10 µl of floral odorant and injected into a constant air stream (1L/min) leading to the moth. The stimulus was pulsed through a solenoid-actuated valve controlled by an RZ2 bioamplifier processor (Tucker-Davis Technologies Inc, Florida). The outlet of the stimulus cartridge was positioned 2 cm from and orthogonal to the center of the antennal flagellum. Stimulus duration was 400ms, and five pulses were separated by either a 5-s interval or 10-s interval. The inter-stimulation duration was approximately 1 min. The tested stimuli were categorized as behavioral and non-behavioral if the mixture contained three compounds: Benzaldehyde (BEA), Benzyl alcohol (BOL), and Linalool (LIN). For our first sets of an experiment, we tested behavioral stimuli including extracts of Datura flowers (DatExt), 5 artificial mixtures (P3, P4, P5, P7, P9) containing the behavioral components, and 3 dilutions of one of either P7 (P7 10, P7 100, P7 1000) or P9 (P9 10, P9 100, P9 10000). The non-behavioral stimuli include mineral oil (control, no odor), 5 mixtures of non-behavioral components (M2, M3, M4, M5, M6), and 9 individual non-behavioral components. We have also presented the moth with P7 without Benzaldehyde. In the second experiment, to determine how modifying the ratio of compounds in the mixture modified the encoding of the floral odor, we used an odor cartridge containing the P7 mixture and a second odor cartridge containing increased concentrations of Benzaldehyde (10-, 100-, or 1000-fold higher concentrations), Benzyl alcohol (10- and 100-higher concentrations), Linalool (10- or 100-fold higher concentrations). The odor from the two odor cartridges was released simultaneously into the airline, allowing them to mix before reaching the flagellum. In this manner, the ratio of compounds in the Datura mixture (P7) could be dynamically altered.

### 2.3 Ensemble antennal lobe recording

The odor-evoked responses of 80 units were obtained from 5 male moths. Recordings were made with 16-channel silicon multielectrode recording (MR) arrays (A4X 4-3 mm-50-177; NeuroNexus Technologies). These probes have four shanks (each of 15 µm in thickness) spaced 125 µm apart, each with four recording sites 50 um apart, and have a surface area of 177 µm^2^. The MR was positioned under visual control with a stereo microscope (Narishige, Japan). As demonstrated in Fig 1A and Fig S1, the four shanks were oriented in a line parallel to the antennal nerve. The MR was advanced slowly through the AL with a micromanipulator (Narishige, Japan) until the uppermost recording sites were just below the surface of the AL. Thus, the four shanks of the MR were recorded from four regions of glomerular neuropil across the AL. Ensemble activity was recorded simultaneously from the 16 channels of the MR array by using TDT amplifiers (Tucker-Davis Technologies Inc, Florida). The recorded signal was digitized at 25 kHz per channel by using synapse software (version 98, Tucker-Davis Technologies Inc, Florida).

**Fig 1.**
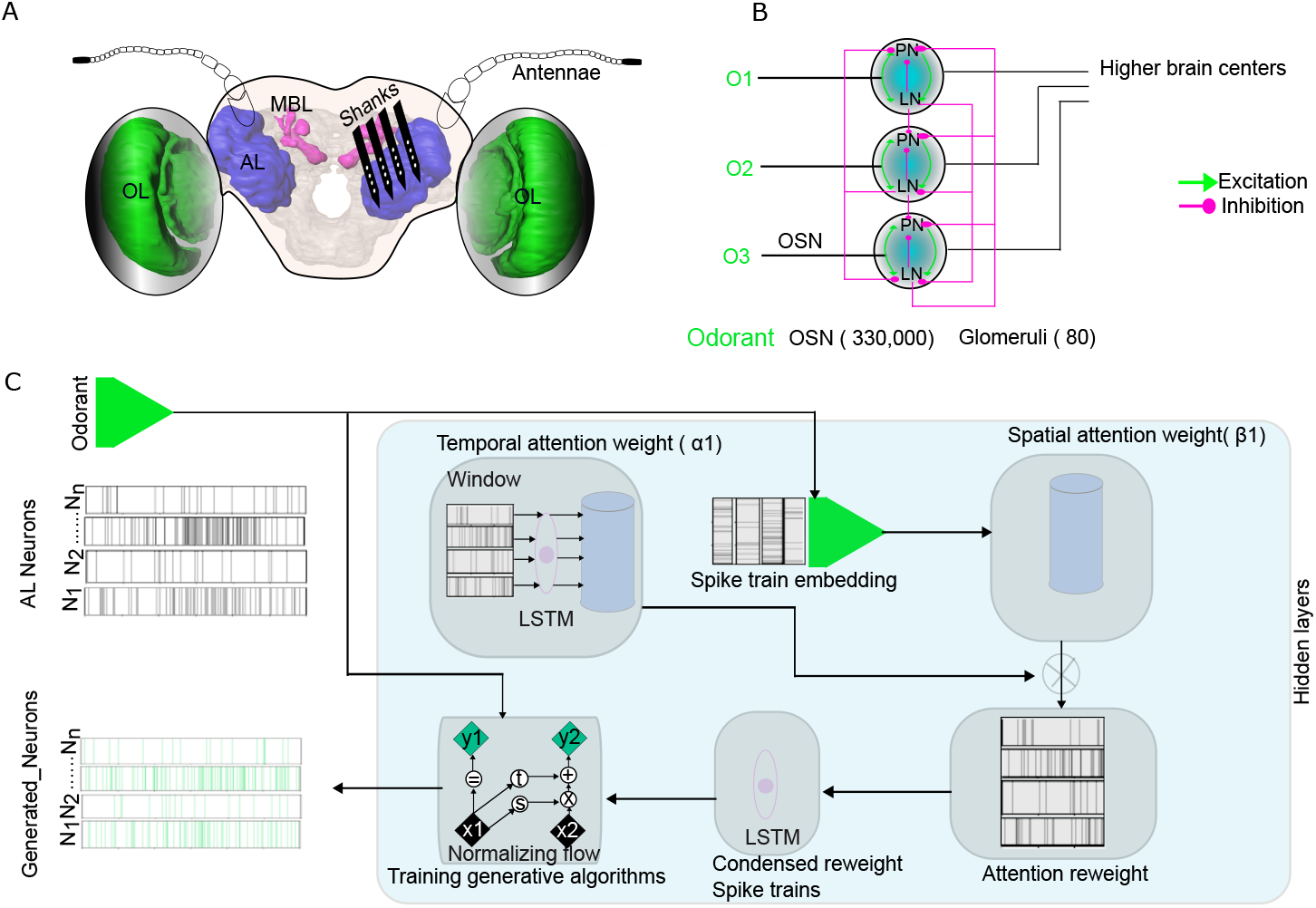
Overview of methods applied in this work. (A) Schematic of the insertion of the probes in the AL. The 3D model of the brain is acquired from (www.insectbraindb.org). Images are not to the scale. (B) Schematic diagram of the antennal lobe network consisting of the projection neurons (PNs) and local interneurons (LNs). The olfactory sensory neurons (OSNs) after activation by the different odorants (O1, O2, and O3), the information is relayed to PNs and LNs. Excitatory connections are denoted by an arrow (green colored) whereas inhibitory connections are denoted by the round head (Magenta colored). The PNs relay information to the higher brain centers. (C) Architecture of the spatial-temporal attention normalizing flow. The spike trains are first passed through a Long short-term memory (LSTM) unit and linear embedding modules to obtain the spatial and temporal attention weights for reweighting the spike train. The reweighted spike train is then passed through a second LSTM module, and its final hidden representation of the reweighted spike train is used as the context vector to train the conditional normalizing flow for learning distributions of the inter-spike intervals (ISIs) and generate realistic spike trains. The *x, y* in the Normalizing flow denotes the input and output of an affine coupling layer [27]; the subscripts 1, 2 different parts of the latent variable; t and s denote two different neural networks. OL: Optic lobe; MBL: Mushroom body lobe.

### 2.4 Localization of recording probes in the AL

The head was excised, and the brains were dissected in the Manduca saline. The brain was washed with 0.01M PBS (2 times: 20 min each) and then submerged in the solution consisting of 4% PFA and 0.03% glutaraldehyde) to facilitate the fixation of the tissue. The preparation was kept overnight at 4^°^C and dehydrated in series of ethanol series (50%, 70%, 90%, 96%, 100%, 100%: 20 min each) and finally cleared in the methyl salicylate (Millipore sigma, USA). The whole mount preparation is scanned with a laser scanning microscope (Nikon, A1R, Nikon Instruments Inc, USA) equipped with a CFI Plan Apo 10X Air objective is scanned with 488 nm line of an argon laser. The high-resolution confocal images with 1024 × 1024 pixels at the distance of 2 – 4 µm in the z – direction were obtained. The image was imported in AMIRA v 6.5.0 (Thermofisher Scientific, USA) and the glomerular structures were reconstructed (Fig S1B). The shank impaled in the AL was also reconstructed and visualized.

### 2.5 Spike Sorting

The continuous waveforms are exported to an offline sorter (Offline sorter, Plexon, Version 4.7.1). The spike data were digitized at 25 kHz per channel. The filter setting (0.6 - 3 kHz and system gain of 1000 were software adjustable on each channel. Spikes were sorted by using a clustering algorithm based on the method of principal components (PCs) (Off-line Sorter; Plexon). Clusters that were separated in 3D space (PC1–PC3) after statistical verification (multivariate ANOVA; P*<*0.05) were selected for further analysis (7-19 units were isolated per ensemble; most units were present in Shank 2 and 3; Fig S1). Each spike in each cluster was time-stamped, and these data were used to create raster plots and calculate peristimulus time histograms (PSTHs). The analyses were performed with Neuroexplorer (Nex Technologies, version 5.4) using a bin width of 5 ms.

## 3 Model

### 3.1 Problem Formulation and Notation

We denote spike train *S*^*q*^ ∈ ℝ^*N×T*^ for a total of *N* neurons, *T* timesteps, and *q* ∈ 1, …, *Q* different stimuli. For the *n*-th neuron, we denote the time of the *i*-th spike timing as 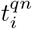, hence the previous spike’s timing as 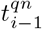. Then the interarrival time between these two spikes is 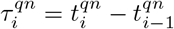. The goal is to model the interarrival time distribution 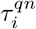 for arbitrary spike *i*.

Let [Δ] denote the window size and denote a windowed spike train as *S*_*i*[Δ]_; we assume the interspike intervals of neuron *n* are conditionally independent given the stimuli *q*, the Δ-windowed history of last spike, and time of the last spike to reduce the problem into modeling the distribution of interarrival time presented in Equation (1). For ease of notation, we drop the superscript *q* and focus on an arbitrary stimulus, we also drop the superscript *n* and focus on an arbitrary neuron.

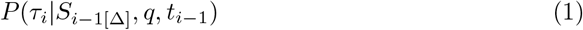

Our generative modeling approach consists of two main components: 1) the spatial-temporal attention units that encode the windowed spike history to latent space; and 2) a conditional generative model learned with a normalizing flow that models the target distribution presented in Equation (1). Beyond the goal of learning the conditional ISI distribution, the attention weights learned by this conditional generative system can be extracted for further analysis. We present the full spatial-temporal attention normalizing flow (STAN-Flow) architecture in Fig 1C.

### 3.2 Spatial-Temporal Attention

The spatial-temporal attention mechanism combines the vision attention mechanism in computer vision [28] and temporal attention in natural language processing [29, 30]. Synchronization can be seen as interactions between neurons which can be characterized through spatial attention, where a higher spatial attention weight corresponds to a higher strength of interaction between neurons. The importance of particular spike timing and the general spiking rate is characterized by the temporal weights that scan through the spiking history: the higher the temporal weight, the more important a specific time is. Therefore, neurons can be synchronized in their activity even if they have different individual temporal dynamics.

The spatial-temporal attention module consists of Long short-term memory (LSTM) layers and a few linear layers. The LSTM layers effectively summarize spike train time series into lower dimensional hidden states, which are then projected by the linear layers to obtain semi-interpretable attention weights. The windowed spike train *S*_*i−*1[Δ]_ is passed through the first LSTM (*f*_1_) and a linear spatial-embedding layer, outputs the hidden representation *h*_*i−*1_ and *d*-dimensional spike train spatial embedding *E*_*i*_ ∈ ℝ^*N×d*^. For the temporal attention, we further reduce *h*_*i−*1_ through a linear layer to obtain the temporal weights, 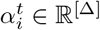.

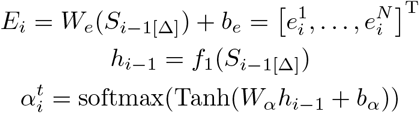

Additionally, Riffell et al. [3] shows that stimuli information is also encoded by the ensemble firing of neurons. Hence we concatenate the last hidden states of the LSTM, denoted as *h*^*^, the spatial embedding of a particular neuron’s activity 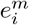, *m* ∈ 1 … *N*, and the stimuli *q* to pass through a linear layer and obtain the spatial weights 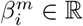. The vector that contains the spatial weights of all neurons is denoted as 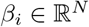. To isolate the higher-order ensemble patterns, we replace the traditional *softmax* of the attention mechanism and apply its sparsifying counterpart *sparsemax* Martins and Astudillo [31] which directly projects logit values onto the simplex. Applying the *sparsemax* allows some attention weights to be reduced to zero, amplifying the effect of those synchronized neurons.

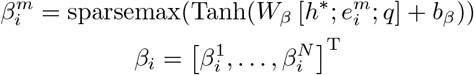

Usual applications of attention weights obtain a context vector through a weighted average of latent variables [29, 30]. However, a weighted average of each neuron’s representation dilutes the synchronization identified through the spatial attention weight as the resulting weighted representation becomes less identifiable. Hence, we reweight the windowed spike train with the mean-normalized spatial-temporal weights and feed the reweighted windowed spike train 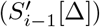 through a second LSTM layer (*f*_2_) to obtain the final hidden representation 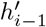. The symbol *×* denotes element-wise multiplication.

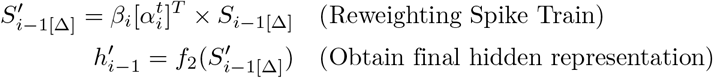

The output of *f*_2_, 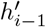, can be seen as a context vector derived from the reweighted spike train. This context vector combines the temporal dynamic spike train, the neuron interaction, and the influence of different stimuli into a continuous representation to inform the conditional generative model.

### 3.3 Conditional Normalizing Flow

Once we learn the synchronization and timing information, we build a modeling module to accurately reflect the ISI distribution based on synchronization and temporal dynamics. While traditionally the modeling of spike train follows the Poisson Process, this assumes the spike train is rate coded and the ISI distribution follows an exponential distribution. These assumptions are not always realistic and constrain the modeling process. Instead of a model based on the Poisson assumption, we build a non-parametric deep generative model conditioned on the final hidden representation 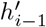 to learn the probability distribution of inter-spike interval given the learned history.

We chose to apply a conditional normalizing flow that directly optimizes the negative log-likelihood of the density. A Normalizing Flow is usually defined by a transformation of a standard Gaussian distribution into a more complex distribution [26]. This transformation normally consists of a sequence of invertible, tractable, and differentiable mappings enabling the evaluation of a sample’s value in the simple distribution or its likelihood.

We concatenate the stimuli *q*, the last hidden representation of attention-reweighted spike train 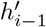, and the time of last spike *t*_*i−*1_ into a context vector denoted as *x*_*i*_. We propose a normalizing flow that is conditioned on *x*_*i*_; the likelihood takes the following form with *Z* being drawn from a conditional Gaussian. Extending recent neural network architecture [26, 27], a loss through log-likelihood can be written as:

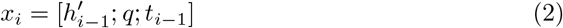

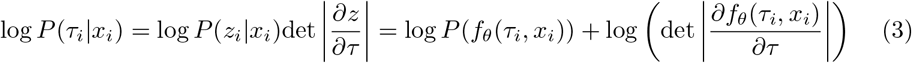

We specifically applied the real-valued non-volume preserving normalizing flow architecture (RealNVP) in our study [27], where the *f*_*θ*_ is characterized through a series of neural networks that construct an upper-triangular Jacobian, simplifying the determinant computation of the Jacobian to be the trace.

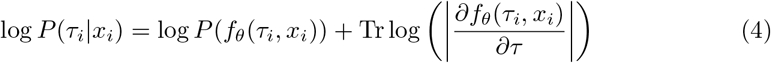

We note that this framework considers the spiking history, the interaction between neurons, and the stimuli effect altogether and learns the ISI distribution without assuming it follows some canonical, parametric distribution. Hyperparameters regarding the architecture and training process are recorded in the Supporting Information.

### 3.4 Identifying Synchronization

A crucial part of our analysis is establishing a higher-order non-linear method to analyze neuron synchronization. We propose the spatial attention weight method and compare it with two previously reported neuron synchronization methods, the Ensemble Synchronization proposed in Lei et al. [32], and the Kernelized Binless Method applied in Martin et al. [14]; we then discuss the synchronization analysis process of our proposed spatial-attention weights method.

#### Ensemble Synchronization

The traditional analysis of ensemble patterns utilizes the cross-correlation coefficient between pairs of neurons Lei et al. [32]. In particular, the synchronization index (SI%) is calculated as

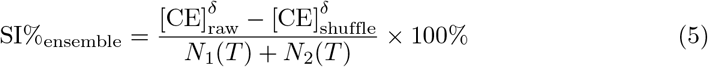

where [CE]_raw_ is the number of coincident events in cross-correlogram peak centered around *t* = 0 with width δ, [CE]_shuffle_ is the number of coincident events after trial shuffling (shift predictor method) to correct for coincidences attributable to chance and an increased firing rate. The corrected correlograms were calculated by averaging four trial shifts and subtracting the result from the raw correlogram. T is the total response time over which spikes were counted, and N1 and N2 are the number of spikes recorded from units 1 and 2 during time T [32].

We calculated the ensemble SI% for all stimuli using one trial as the raw trial and corrected with shuffling the other 4 trials. We applied the parameters δ = 5, and *T* = 1000 (msec) after the onset of the stimuli as suggested in [3]. In the supporting informations (Fig S2), we explore a variety of hyperparameters for δ and *T*.

#### Kernelized Binless Method

A more recent method that analyzes the synchronization of neuron firing is through the kernelized binless method [14]. While it remains a pairwise synchronization analysis, it applies an exponential function kernel to smooth out the spike train. Specifically, the exponential kernel is denoted *h*(*t*) = exp(*−t/τ*)*/u*(*t*) where *u*(*t*) is the heavy side function, *τ* is a kernel parameter to aggregate spikes over time; a similarity index (see Equation (6)) is then calculated between a pair of neurons’ kernelized spiking.

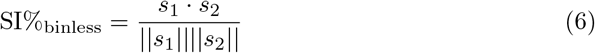

We compute SI%_binless_ with the time constant *τ* = 5(msec) similar to previous synchronization analysis [14]. Trial shuffling is also applied for the kernelized binless method. In the supporting informations (Fig S3), we explore a variety of hyperparameters for *τ*.

#### Spatial Attention Method

We train a conditional normalizing flow for each neuron by applying a cross-validation scheme in which we rotate 3 trials to form the training set while the other two form the validation and test set. The spatial attention module (see Section 3.2) are learned jointly with the conditional normalizing flow through the loss function (4). For an arbitrary spike *i* of an arbitrary neuron *n*, and arbitrary stimuli *q*, our modeling process would determine a set of attention weights that determines the importance of each neuron in the neuron population. During our analysis, we concatenate the spatial attention for each stimulus, then average the spatial attention weights over all spikes, all neurons, and all runs during evaluation to output a synchronization summary matrix *B*. The specific calculation is shown in Equation (9).

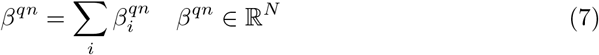

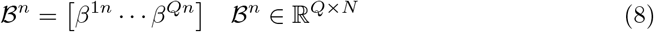

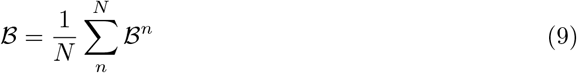

### 3.5 Classification of neurons into putative PN and LN

Although our ensemble recorded neuronal data does not allow us to identify the neuron types, we follow the classification procedure described in Lei et al. [33] to classify PNs and LNs in our spike-sorted units. This classification method relies on the observation that the spontaneous spiking activity of PNs and LNs is different: PNs are more likely to have bursts of spiking activity while the LNs fire regularly. It adopts the criterion in Legendy and Salcman [34] to detect potential bursts in spontaneous activities (5s) in the full spike train from Poisson Surprise (*S*) rates, which characterizes the abrupt changes spiking rates compared to the mean spike rate.

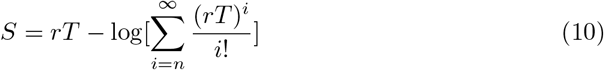

The Poisson Surprise rate for a set of spikes is computed from the time span *T* of the set and the mean firing rate *r* which is the number of spikes *n* in the set divided by *T*. The burst is detected by first finding a pair of successive spikes whose inter-spike interval (ISI) is less than the mean ISI of the spike train multiplied by a designated coefficient *p*(0 *< p <* 1). Subsequent spikes are added to the pair of spikes to formulate a spike set with the largest possible Poisson Surprise value, and the earliest spikes are pruned from the set if that further increases the Poisson Surprise of the spike set. Finally, the spike set is regarded as a burst if it consists of at least 3 spikes.

With all burst occurrences detected throughout the spike train, we use them to calculate 9 burst-related features for a particular neuron Lei et al. [33]. A logistic regression is finally fitted with the 9 burst-related features as covariates to classify the type of neurons as PN or LN.

We train a similar logistic regression based on the spontaneous spike train obtained from intracellular recordings and stainings in Lei et al. [33], the validation accuracy is around 85%. Then using this logistic regression model, we classify the neurons collected through section 2. During our initial data analysis, we found that the distribution of the average spike rate of the data from [33] is different from our spike-sorted data. The difference in distribution resulted in scale differences in the 9 burst-related features. To resolve the difference in the features, we applied the following processing steps:

1. We tune the *p* parameter in the procedure for detecting potential bursts to obtain burst-related features in a similar scale. We used *p* = 0.2 while *p* = 0.5 is defaulted in Lei et al. [33]. The *p* parameter defines the ratio between the mean spike rate (*mr*) and the spike rate of potential burst segments (*br*) and classifies the segment as burst when *mr/br < p*.
2. We remove three of the 9 features where the significant scale differences cannot be resolved by the tuning of *p*. The 6 features we used to classify the neuron types are within-burst max spiking frequency, within-burst number of spikes, percentage of burst spikes, burst frequency, mean Poisson surprise, and max Poisson surprise.
3. We apply two different min-max scalers to the training data [33] and the testing data (described in section 2), respectively.

Once the neurons are classified, we supply our predicted labels to human experts to assist in the annotation of true neuron types. We refer the readers to Lei et al. [33] for the details regarding the classification method.

### 3.6 Response index and Cosine similarity

#### Response Index

Response Index was computed in our study to investigate the response of different units to the odor stimuli and also to assess the similarity of generated and real spike trains under different stimuli for every units (Riffell et al., 2009). The response index is calculated as the following:

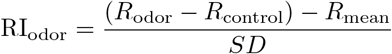

where the *R*_odor_ is the response of a specific odor; *R*_control_ is the response of control (mineral oil); *R*_mean_ is the mean response averaged over all stimuli, and *SD* is the standard deviation of the response across all stimuli. Response in this study means the number of spikes over the stimulation period (0-400ms after the onset of stimuli).

#### Cosine Similarity

Cosine similarity was used to study the similarity between synchronization patterns of different stimuli. The cosine similarity essentially measures the angle between two vectors; the values of cosine similarity reside in [*−*1, 1] for −1 being the least similar and 1 being the most similar. For any two vectors *v*_1_, *v*_2_, the cosine similarity is defined as:

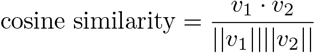

The denotes the dot product, || · || denotes the norm of a vector.

## 4 Results

By leveraging known biological phenomena of the spiking and interaction between neurons in the primary olfactory center, the antennal lobe (AL) acquired through inserting silicon multielectrode array in *Manduca sexta* (Fig 1A and Fig 2B), we designed STAN-Flow to model the fundamental neuron spiking mechanism and neuron interactions in the AL (Fig 1C). We validate our method from three different perspectives: 1) can STAN-Flow generate realistic spike trains that follow the distribution of collected spike trains? 2) can STAN-Flow learn to discriminate behavioral and non-behavioral stimuli through its spatial attention weights that mimic the synchronization of neurons? 3) can STAN-Flow automatically detect the interactions between different types of neurons in the AL region? As we find convincing evidence that SPAN-Flow indeed learns the neuronal spiking and interaction activities, we then use STAN-Flow to test whether the pattern of synchronization between PNs and LNs is reduced when the component odorants differ from their concentrations in behaviorally relevant complex odor mixtures (e.g. floral scents that trigger foraging). The related formulas used for evaluations including the Response Index (RI) and cosine similarity are described in Supporting Information A.

**Fig 2.**
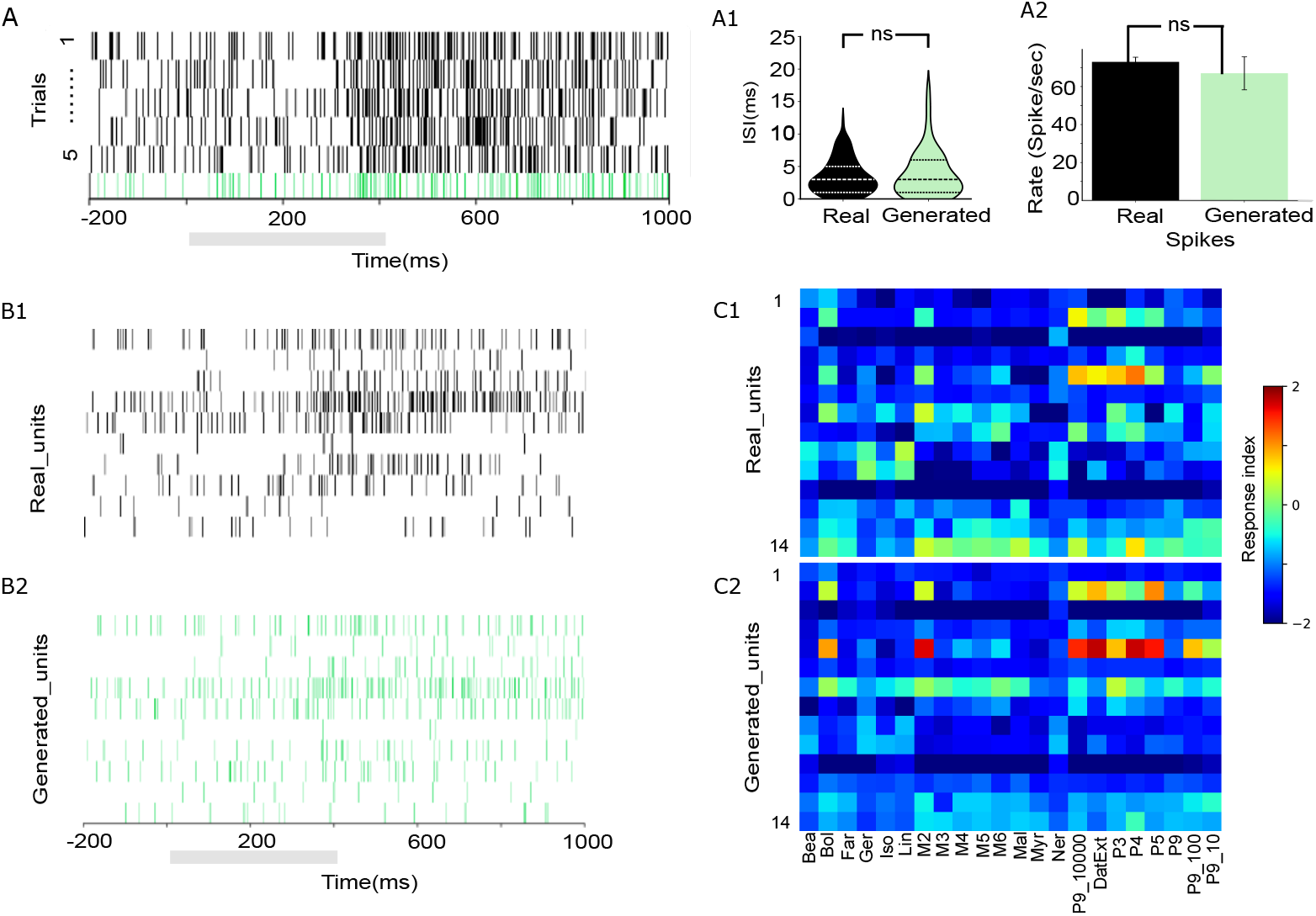
STAN-flow learns the generating distribution of ISI. (A) Comparison of Real spike train of a particular projection neuron across 5 trials (black) vs a generated spike train of the same neuron (green) under P9 stimuli; the grey bar indicates stimulation window. (A1) Violin plot of ISI of real and generated spike train in (A) under P9 stimuli the dashed line indicates the 25%, 50%, and 75% quartile. (A2) Comparison of real vs generated spikes’ average spike rate. (B1) Real spike trains of 10 neurons stimulated with P9. (B2) Generated spike trains of 10 neurons stimulated with P9; the grey bar indicates the stimulation window. (C1) Response Index calculated with a population of neural response to the floral scents. (C2) Response Index calculated with model generated population neural response as (C1). ns: Not significant

### 4.1 Antennal lobe network dynamics and spike train generations

One major aspect of validating the STAN-Flow architecture is examining how similar the generated spike trains are compared to the real neuron recordings. In Fig 2A, we present the spike train of a specific unit in response to an odor with 5 trials. We note that odor-evoked activities start about 400 msec after odor stimulation. At around 400 msec the neuron starts a burst of activity from the stimulation, and then the spiking activity diminishes after 800 msec. The generated spike train (green, Fig 2A) realistically captures this change in firing rate before, during, and after the stimuli. We conclude that there are no differences in the real and generated distribution of ISI (Kolmogorov–Smirnov (KS) Test^1^, P *>* 0.05; Two One-Sided T-Test (TOST) for distribution mean^2^ [35], P*<* 0.05). We also found no significant differences in the average firing rate of a neuron for real and generated spike trains (two-sided, two-sample t-test, df=8, P*>*0.05), indicating that the generated spikes are similar to the real ones (see Fig 2A1 and Fig 2A2).

We also examine the generated spike trains across a neuron population. We present the real and generated spike trains of 10 units stimulated with a behavioral odor, respectively (Fig 2B1 and Fig 2B2), showing that STAN-Flow generated spike train changes correspond to the temporal dynamics of the real neural response. We compute the Response Index (RI; Fig S4 according to the computation formula described in Supporting Information A) across all neurons and stimuli. The ensemble responses to the floral scent for the generated spike trains correspond well with the real indicating that the STAN-Flow follows the spiking pattern of the actual recordings across different stimuli and neurons. This multi-neuron and multi-stimuli comparison further confirms the state-of-the-art modeling capabilities of STAN-Flow in learning the spiking process of neurons in the AL region.

### 4.2 Spatial attention weights classifies stimuli

The AL relies on different cell pairs synchronizing with one another, or a specific subset of the critical neurons to encode the behavioral and non-behavioral stimuli [3, 36]. Our method capture this synchronization mechanism through the spatial attention module.

The spatial attention module assigns an attention weight to each of the units of the spike train input. The attention weights are then rescaled by the mean to reweight each unit’s spike train for generative modeling. We extract the spatial attention weights and compute a summary matrix *B* through Equation 9, where each row of *B* represents a stimulus, and the columns represent the relative importance of each unit and hence is a measure of synchronization that includes activity distribute across time and across neurons with different individual spiking dynamics. The spatial attention method takes all neurons into account during the modeling of spike trains, thus offering the ability to characterize the synchronization of multiple neurons beyond pairwise analyses.

We apply t-distributed Stochastic Neighbor Embedding (TSNE) to compare our method with other synchronization analysis methods [37]. We compute the pair-wise synchronization matrix for pair-wise methods including the Ensemble Synchronization Index [3] and the Binless method [14]. Then through TSNE, we reduce the dimension of *B*, and the upper-triangular matrix of the pair-wise synchronization method of Ensemble method [3] and Binless method [14] into two dimensions and present the 2D scatter plot in Fig 3. Our result demonstrates that the spatial attention weights distinctly separate the stimuli into two clusters: one includes all the stimuli mixtures that contain all the behavioral components, and the other cluster includes the individual odor molecules as well as mixtures containing non-behavioral components (Fig 3A and Fig S5). Two exceptions are grouped with the behavioral relevant stimuli space: the single odor benzyl alcohol (BOL) and M2 (mixture containing BEA and BOL: Riffell et al. [3]). These results suggest that BEA and BOL could be essential to the behavioral responsiveness of complex odors. In addition, we compare the Ensemble synchronization index (ESI) and Kernelised binless (KB) method performed on our datasets that fail to display any such clustering effect(Fig 3B and Fig 3C). We measured classification accuracy through a 2-class K-means algorithm to test the methods’ ability to separate the behavioral and non-behavioral stimuli and found that our method is better at classifying the oder by their functional relevance. While there is no significant difference between the Ensemble synchronization and the Binless method with an accuracy of around 60%, our method offers an enhanced accuracy of around 80%. We also note that we iterated our method 100 times with random initialization, and it consistently outperforms the other methods (One-sided z-test, Ensemble Synchronization Index: p-value*<*0.01, Binless method: p-value*<*0.01). This enhanced result shows convincing evidence that spatial attention is a more robust method for characterizing synchronization in a complex setting. The other methods were previously only applied to populations of projection neurons [3, 14]; we conjectured that ESI and KB failed to separate the stimuli types in this neural population because both LNs and PNs are present. This enhanced performance in separating the behavioral and non-behavioral stimuli when different types of neurons exist in the neuron population propels us to understand how the spatial attention models the interaction between LNs and PNs.

**Fig 3.**
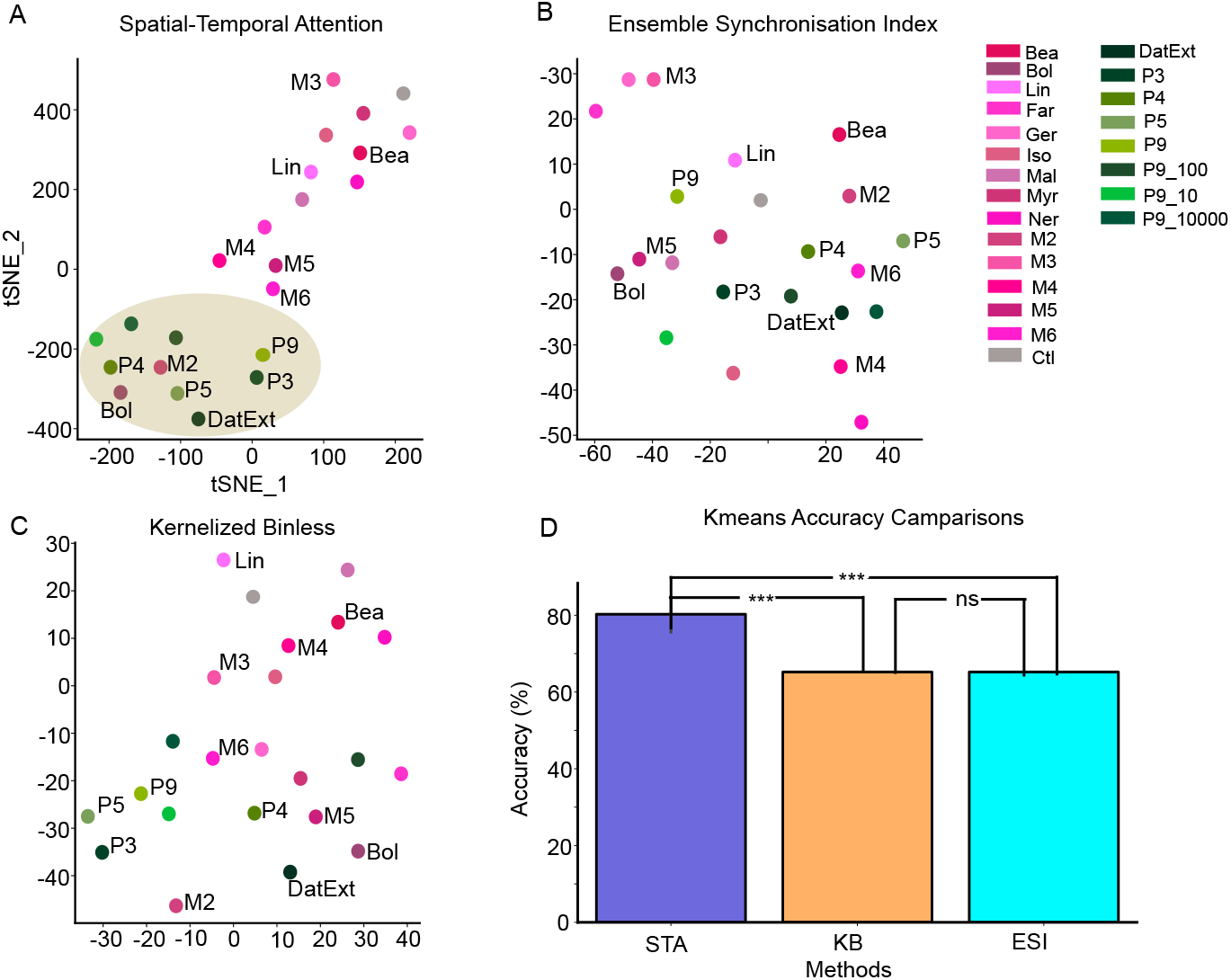
Clustering behavioral and non-behavioral stimuli through different synchronization methods. (A) Spatial attention weights separate the behavioral stimuli (represented by different shades of green) from the non-behavioral stimuli (different shades of Magenta) by forming two distinctive clusters in the 2D TSNE reduced space. The behavioral stimuli cluster is highlighted with a brown background. (B) Elements of the upper triangular matrix are extracted as a vector from the pair-wise Ensemble Synchronization Index matrix, and then reduced to 2D through TSNE. (C) Similar to (B), but we reduced the upper triangular matrix of the pair-wise Binless synchronization through 2D TSNE. (D) Bargraph represents the 2-class Kmeans algorithm with the Spatial-temporal attention (STA) method repeated with 100 different initializations. Kmeans accuracies of the Ensemble Synchronization index (EI, P*>*0.05) and Kernel Binless (KB, P*>*0.05) clustering are significantly less accuracies to STA. *** denotes P*<*0.001 and ns is non-significant

### 4.3 Detecting the interaction between PNs and LNs

We have shown that the spatial attention is robust in dealing with multiple neuron types, allowing us to capture the synchronization mechanism that separates different stimuli. We now analyze the interaction relationship between different types of neurons through spatial attention. Our multi-unit recording in the AL, however, does not permit a morphological identification of the PNs and LNs. To classify the neurons, we utilize previous results that found the PNs and LNs have distinctively different spontaneous patterns of spiking activities (Fig 4A) [33]. The PNs burst from time to time during spontaneous firing, while the LNs spike regularly. Based on this observation Lei et al. [33] developed a simple method to classify the neurons based on their spiking dynamics. We apply the method developed in Lei et al. [33] to classify the neurons based on their spontaneous spiking. Instead of the 9 parameters, we measured the six parameters describing the burstiness of the PNs and LNs and fed to the logical regression and categorizing the neurons (we visualize 3 in Fig 4B). We refer readers of the details of this classification process to section 3.5. We then supply our classification result from logistic regression to human experts to further identify neurons with high classification uncertainty. The classification accuracy computed against expert label was 78% while the validation accuracy using data from Lei et al. [33] is about 75%.

**Fig 4.**
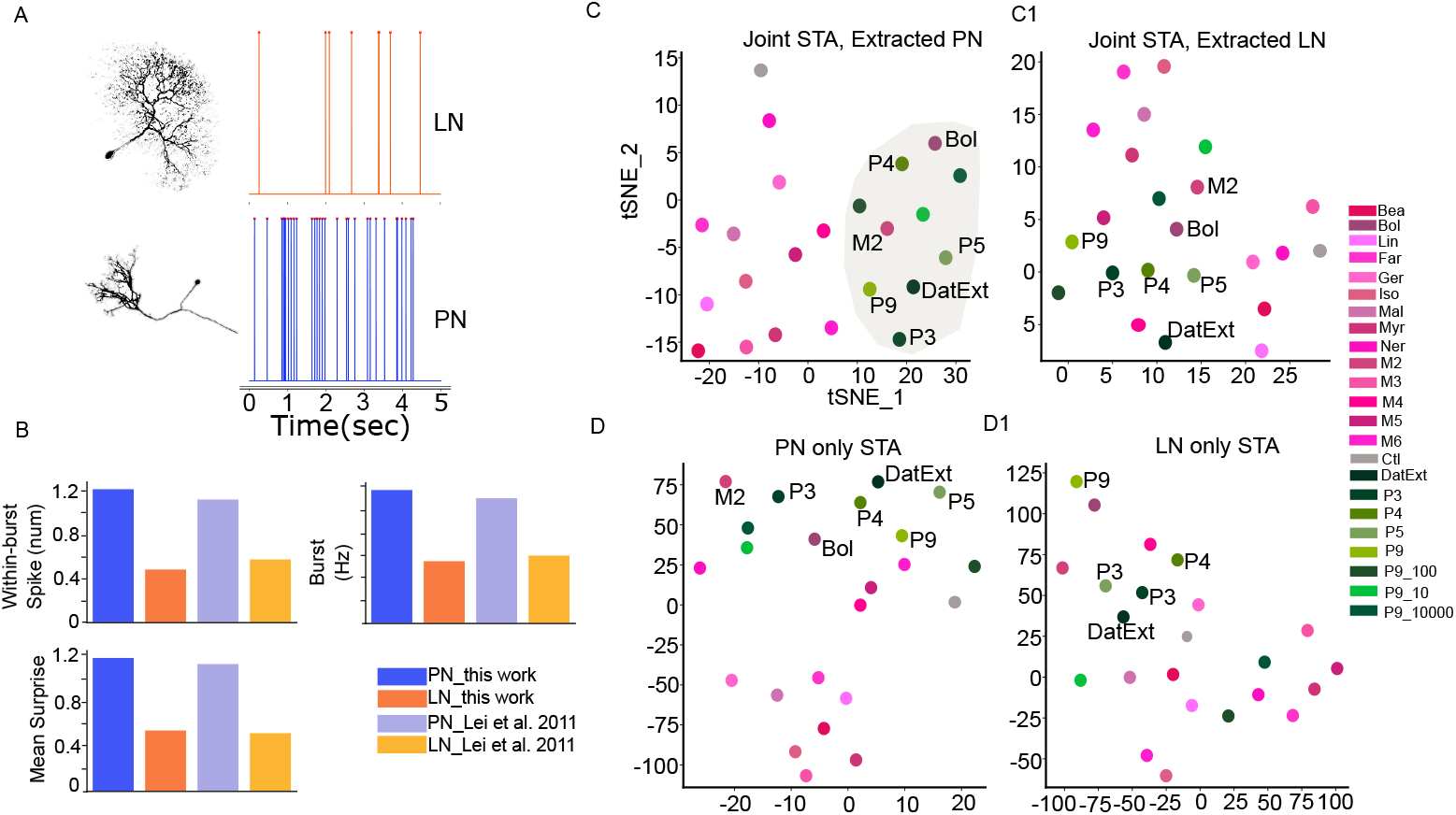
Spatial Attention module learns the interaction between two neuron populations in the antennal lobe. (A) Representative morphology and physiology of the local interneurons (LNs) and Projection neurons (PNs). LN spikes more regularly while PN spikes in a burst pattern. (B) Selected features used to classify neuron types; the difference between PN and LN is consistent with previous analysis of Lei et al. [33]. (C) The spatial attention of PNs and LNs (C1) was separately extracted from a joint model (trained with both PNs and LNs) and then reduced to 2 dimensions through TSNE. The behavioral cluster of stimuli is highlighted with light-shaded gray. (D) TSNE reduced 2-dimensional scatter plot of Spatial attention with the model trained with PNs only and with LNs only (D1).

The essential question in this analysis is to understand the interaction between the PNs and LNs. As Lei et al. [32, 36] and Tanaka et al. [38] suggested, LNs modulate the PNs synchronization, and the PNs synchronize among themselves [14] to encode behavioral and non-behavioral stimuli. We extract the synchronization patterns of PNs and LNs from the joint model, where we train using the entire population of neurons in our recordings, and apply the 2D TSNE analysis to the synchronization matrices.

Results showed that the PNs’ synchronization matrix continued to separate the behavioral and non-behavioral stimuli in 2D reduced space while there were no visible clusters in LNs’ synchronization responses (Fig 4C and Fig 4D). Such behavior, however, is not observed when we trained a STAN-Flow for each neuron type: the spatial attention weights do not cluster for either the PN-only or the LN-only model (Fig 4D and Fig 4D1). The contrasting clustering behavior in Fig 4 suggests that we are able to recover the interaction mechanism between PNs and LNs. To obtain clusterable representation with PNs, LNs must be available during the modeling process–the AL region and the STAN-Flow need the LNs to modulate the synchronization of PNs for properly identifying the behavioral and non-behavioral stimuli where the LNs might inhibit the neuron processing the non-behavioral component (Fig 4C and Fig 4D). Together, these results highlight the functional significance of the LNs in PNs’ synchronization.

Our previous result has highlighted that LNs are critical for odor classification, but could there be a core neuronal unit in an ensemble that accounts for the segregation of behavioral and non-behavioral stimuli? As we showed that spatial attention consistently clusters the behavioral and non-behavioral odors, we now test the Kmeans clustering accuracy by removing each unit in an ensemble. We found one LN significantly lower the classification accuracy (86% before removal vs 78% after removal). This neuron responds to both behavioral compounds and non-behavioral compounds Fig 2C1, unit 14). In addition to single neuron analyses, we removed a combination of up to 3 neurons (data not shown). We found a specific combination including the LN (unit 14, Fig 2C1) and two other PNs (units 2 and 5, Fig 2C1) will lower the clustering accuracy to 52%. These PNs responded to most of the behavioral compounds. These results further indicate that broadly tuned LNs could be core neurons for odor classification in combination with the PNs which are responsive to the behavioral compound only.

### 4.4 Altering the excitatory drive attenuates synchronization

The previous analyses showcase STAN-Flow’s accurate recovery of the general functionality and spiking activity of the AL region. We showed that STAN-Flow can not only generate realistic spike trains for different neurons and different stimuli but is also able to discriminate behavioral and non-behavioral stimuli through its spatial attention module that mimics the neuron synchronization of the AL region. We also show that the attentional method can recapitulate the interaction relationship between PNs and LNs by examining the synchronization pattern of each neuron type, even demonstrating that the clustering of odorants into behaviorally relevant stimuli occurs in the PNs but is dependent on underlying LN dynamics. Knowing that STAN-Flow captures the synchronization dynamics of the AL, we next tested the hypothesis of how the neural synchronization and clustering are tuned to fluctuating naturally occurring odorant concentrations. We changed the odorant ratio by altering the component concentration of benzaldehyde (BEA) in a behaviorally relevant odor mixture (P7). We retrained STAN-Flow with neural activities from individual compounds, P7, and P7 with added BEA (10, 100, and 1000 fold increase). We first repeat the TSNE analysis with the summary matrix *B* (see Fig 5A). This module clusters the individual components to the top right corner, the P7 with different BEA intensities in the middle, and control to the far left.

**Fig 5.**
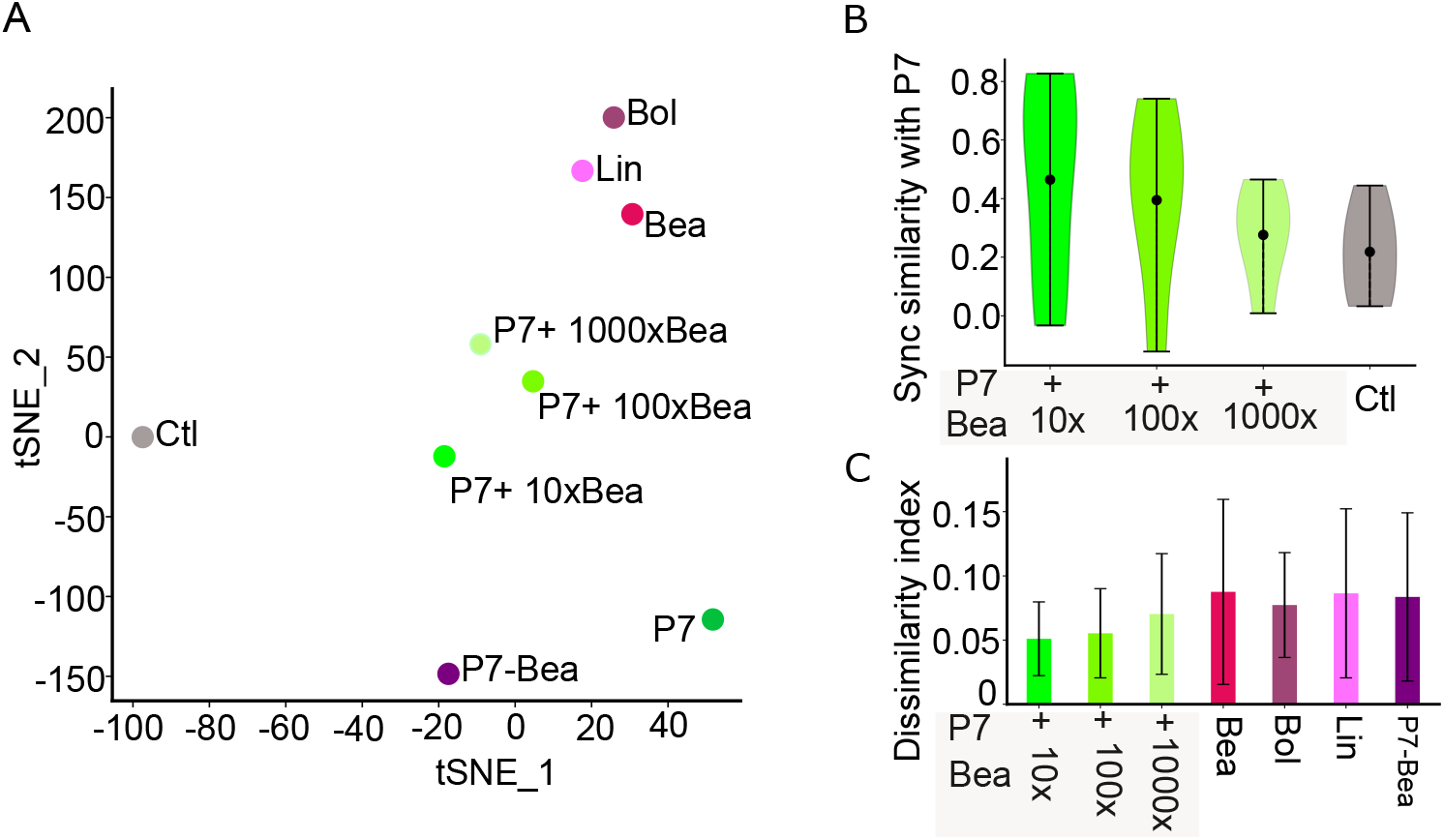
Changes in synchronization pattern with increased Bea ratio in P7. (A) Two-dimensional scatterplot of the TSNE reduced spatial attention weights. (B) Cosine similarity of synchronization pattern between P7 and P7 with increased Bea concentration in the mixture (n = 5 trials). (C) Euclidean distance of P7 spatial attention weights computed against P7 with increased Bea ratio, individual behavioral component, and P7 without Bea.

To compare the ensemble representation of the increased concentration of BEA, we compute two different metrics to analyze the similarity and dissimilarity of synchronization patterns. We chose to apply cosine similarity (see Supporting Information A) that measures the similarity patterns between the P7 mixture with increased BEA ratio and the original P7 mixture. Interestingly, the similarity decreased linearly from a 10-fold increase to a 1000-fold increase in the concentrations of BEA (Fig 5B). The second analysis examined the odor-evoked responses of the increased concentration of BEA in the multivariate space through normalized Euclidean distances (normalized by mean). We compute the Euclidean distance (dissimilarity index) between the P7 odors and different intensities of BEA and also the individual odor compounds. In Fig 5C, We found that the higher concentrations of BEA in the P7 mixture induced a more dissimilar synchronization pattern to those of P7 indicating that the increased concentration of BEA could lead the moth to identify these stimuli as non-behavorial. Our finding further indicates that the neurons in the AL region use synchronization to encode and classify different odors. The neural synchrony is sensitive to changes in the compound concentration or its proportion in the odor, leading to their efficient identification of behavioral odors under various chemical stimulus conditions in the wild.

## 5 Discussion

Here in the current study, we elucidated the effects of network dynamics in the antennal lobe (AL) to classify behavioral and non-behavioral relevant odors. We developed the spatial-temporal attention normalizing flow (STAN-Flow), an accurate computational model representing the spiking ensemble dynamics of the AL. We adopt this model to extend the characterization of the AL network beyond the experimental possibilities. The model effectively reproduced the key features of AL responses in relation to the odor classification. This model also demonstrated that local interneurons play critical roles in the temporal encoding of odor stimuli, enabling the classification of odors into behavioral and non-behavioral stimuli. Shifting the concentration of one of the behavioral compounds in the odor mixture - by altering the balance of excitation and inhibition in the AL - causes the neuronal representation of this stimulus to change. This computational model can be easily modified to be applied in various fields for accurately modeling and reliably interpreting complex interactions for biological and non-biological systems.

### Neurophysiological computation in the AL

The AL is one of the most studied neural structures of the insect brain in terms of detailed cellular- and circuit-level architectures for sensory encoding. Over the last two decades, behavioral, physiological, and modeling research has made great strides towards understanding the circuit basis of processing complex odor mixtures, their intensities, and also their relation to odor classification. Understanding the role of the AL in odor perception has been the focus of a variety of theoretical and computational models. The dynamic and complex stimuli necessitate utilizing the computational models to extract the features of interest from the spike trains [22]. The STAN-Flow developed in this study serves this purpose: its flexibility simulates the spike train generation process and successfully discriminates and classifies complex odors. This model could be beneficial in identifying future odors whether or not they could be relevant to insects, predicting the population response, and simulating the spike trains of the neurons.

This computational model can cluster the odors into behavioral relevant and non-behavioral relevant groups (Fig 3 and 5). Our approach improves clustering into behavioral and non-behavioral odor stimuli and is highly efficient. It could potentially facilitate further processing in the higher brain centers such as the lateral horn (LH) [39, 40]. Given the ample neuromorphic knowledge on downstream neurons from the AL and its circuits, it is still unknown, to date, how this spatio-temporal information is processed within the AL and in higher brain centers.

There remain a few limitations in the STAN-Flow architectures and related analysis. One main limitation is the interpretability of the neural networks. Although, through post-hoc analysis, we showed that the spatial attention weights can be interpreted to a great extent, there are limited theoretical analyses on the attention module to guarantee the interpretability of the attention weights. There are, however, ways to improve the credibility of the result: one can cross-validated with established results to check if the interpretability matches expectations, as we did with PN and LN classification. In addition, one could repeat the experiments many times (e.g., 100 different initializations) and across subjects to verify the consistency of the result. Another drawback of this current architecture is its scalability. The current training scheme of STAN-Flow for performing the group-wise synchronization analysis requires neuron-specific STAN-Flow; while the number of models scales linearly with the number of interacting neurons, it remains difficult to analyze for a larger population of neurons at a greater scale. Given different modeling scenarios, however, this could be resolved by a more intricate STAN-Flow that can be modified to model synchronization hierarchically when dealing with neurons from different regions; it could also be easily adapted to model a multi-dimensional time series instead of modeling each neuron at a time. Finally, the recent rise of diffusion generative modeling techniques can be applied to improve the normalizing flow [25, 41]. There also exist ways to directly connect the interaction between neurons with the generative component. For example, the interaction between neurons can be connected to interacting particle systems that are characterized through a Mckean-Vlasov diffusion, which can be applied as a latent process to greatly improve the interpretability of the machine learning system [42].

#### Local interneurons necessitate the PNs synchronous activity

The olfactory information is encoded as spatial-temporal patterns in the neural population in the AL, and through the activity of different cell types, such as LNs and PNs. A benefit of the model and resulting analyses provides a dissection of the contribution of different cell types in how the complex odor stimuli are processed. However, the diversities of the PNs following the different antennal lobe tract are not considered due to unavailability of the neuroanatomical data following recording. From this study, recorded neurons in the AL are two putative types: Projection neurons (PNs) and local interneurons(LNs). Most PN responses are dynamic under different odors; some are activated, and some are inhibited with no responses (Fig 2C, Fig S6) due to the interaction of the olfactory sensory neurons (OSNs) and LNs. The excitatory feedforward information from OSNs activates PNs, and the PNs also will also indirectly activate the LNs. It has been speculated that a small subset of glomeruli and associated PNs may be required for the valence-specific response (behavior or non-behavior). Here these neurons could typically exhibit stable and synchronized activity to the odorant mixture even though there is minor perturbation in environment [43–46]. Most of the studies done previously are tested on the binary mixture and a few studies have considered more than two compounds in the mixture [3, 47].

The inhibitory LN connecting the defined subsets of the glomeruli could play a crucial role in understanding the perceptual constancy in the olfactory circuits, especially in understanding the synchronized activity of the PNs to the behaviorally relevant mixture. Previous work has shown pharmacological receptor antagonists, targeting the GABA receptors, abolished the synchronized activity of AL neurons and affected the olfactory behavior of the moths [5, 33]. The LNs modulate the temporal patterns of PNs spiking responses, resulting in odor-evoked activity which can enhance the synchrony of sister PNs within the same glomerulus as well as the synchrony of co-activated PNs from the other glomeruli [13, 14]. However, the structural connectivity information is still lacking in *Manduca sexta*, which limits our understanding of the synaptic-level connections between neurons. Electron microscopical studies in fruit flies have shown that the OSNs contribute 75% of the synaptic input to PNs and the remaining 25% is contributed by the LNs [48]. The local interneurons have diverse and complicated spiking patterns [18, 49]. It also has different innervation patterns (within the glomerulus vs across the glomerulus) and innervation targets. It is possible that an LN that makes specific synaptic connections with a given glomeruli could provide the postsynaptic inhibition to the PNs that are processing the non-behavioral relevant odors and receiving information from the subset of behaviorally relevant neurons. This can be ecologically relevant to inhibit the input of the non-behavioral stimulus pathway since they might be the background. In a variety of organisms such as moths and fruit flies, the LNs contain both pre and postsynaptic synapses, and the distribution density of these synapses is biased across different glomeruli, and this bias could eventually affect the extent of the lateral inhibition processing mixture [50, 51].

#### Concentration varying effects on AL network dynamics

Navigating through the complex and dynamic olfactory environment, the moth is challenged with fluctuating odor concentrations. The moth should evaluate the odor and also its intensity. Behaviorally, *Manduca* has been shown that a subset of the behaviorally relevant odorants is processed in a quick (*<*500 ms) and reliable manner [4]. Suppose one of these compounds is removed in the 7-component behavioral mixture. In that case, they are evaluated as non-behavioral and the neural population responses are clustered outside the neighborhood of the 7 component floral behavioral mixture (Fig 3A and Fig 5A) [3, 5]. The behavioral compound such as BEA could play a significant role in the discrimination of the odors in behavioral or non-behavioral and thus affect the olfactory navigation.

The decreasing intensity of the floral mixture, diluted up to 1000-fold clustered with the Datura floral mixture (This study, Fig 3, [3]) could be due to gain control of the LNs [52]. Altering the ratio of Benzaldehyde in the floral mixture, we noticed that different ratios are clustered outside the neighborhood of the floral mixture. The concentration of the same odor may have different or even opposite values [44] and this odor could modify the quality of the odor [53]. We suggest that the excitation and inhibition drive of the local interneurons plays a crucial role in such clustering when the ratios of compounds are altered [54]. After pooling the PNs and LNs separately, we were surprised that altered ratios were separated from the floral mixture, but examining the LNs only, the floral mixture was clustered together with the floral mixture containing a 10-fold high concentration of BEA (Fig S6). One potential hypothesis is that the increased proportion of BEA in the mixture suppresses the PNs that encode the BOL and LIN components, and this will alter the excitation/inhibition balance and decrease the synchronized neural activity. It is also likely that altering other behavioral components could induce greater suppression.

#### Significance and Broader Impact

Our STAN-Flow explores a new, computational avenue to study interactions within neural systems. The interaction of multiple sensory systems is common in many biological organisms allowing these organisms to respond swiftly and efficiently to complex and dynamic environments. For instance, in other animals and humans, multiple peripheral, central, and motor systems work in concert to produce coordinated behaviors. One prime application of the spatial-temporal attention module, which is a significant focus of our ongoing research, is the modeling of multisensory binding in the brain that is funneled downstream via the descending neurons to the motor program. The neural mechanism of integration of multisensory information in the brain is not known and how they drive the motor program is still in investigation. The motor programs involve the coordinated activity of individuals and groups of flight muscles that interact dynamically to produce agile movements and abrupt changes in behavior [55]. Despite extensive research, the interactive mechanisms that govern these muscle dynamics remain largely unknown [56]. The spatial-temporal attention module has the potential to uncover these mechanisms by providing a framework that captures the intricate timing and spatial relationships involved in motor coordination.

Another significant benefit of the STAN-Flow model is its deep generative component. Biological interactions are inherently stochastic and often do not conform to traditional statistical distributions such as the Poisson or Gaussian distributions [57, 58]. This stochastic nature presents challenges for conventional modeling approaches that rely on these distributions. The introduction of a flexible generative model through neural networks, as seen in STAN-Flow, allows for more accurate modeling of natural phenomena with fewer assumptions about the underlying distributions. This flexibility is particularly important when exploring complex biological interactions that may be high-dimensional and highly non-linear. By leveraging the power of neural networks, STAN-Flow can capture the rich, varied nature of biological data and provide deeper insights into the interactions between multiple brain regions or systems.

STAN-Flow’s success in modeling the antennal lobe region has illuminated the potential of combining attention mechanisms with deep generative neural networks to understand the complex interactive relationships between organisms and their environments. The ability of STAN-Flow to accurately model the dynamic and non-linear interactions in the AL region suggests that similar approaches could be applied to both biological and non-biological systems. This attention module could be applied to other sensory systems such as the vision and the auditory system for stimuli discrimination[59]. However, whether one of the systems favors one of the modules (spatial or temporal) is elusive. This opens up new avenues for research in understanding how different neural systems interact and adapt to their environments, ultimately contributing to a more comprehensive understanding of biological complexity and adaptability. Apart from the biological context, STAN-Flow can be used in the digital field [60]. The STAN-Flow can be naturally applied to related tasks such as analyzing videos and speech by summarizing the interaction of different graphic regions and condensing the importance of different periods of videos. By enforcing a set of constraints on the attention weights, it is also possible to extend the module to track objects through space and time. As many climatic phenomena also originate from interactions of local climate [61], STAN-Flow also provides a generative predictive algorithm that allows explicit interaction between local climates to forecast weathering trends in the future. In general, the flexible, semi-interpretable neural network structure of STAN-Flow offers a wide range of applications that can help inform scientists with high orders of interaction between groups or individuals and through time.

## Author Contribution

HY, PKC, VT, and JR conceptualized the study. Data curation was carried out by JR, PKC and HL, while HY, PKC, and PC conducted the formal analysis. Funding for the research was secured by SS, VT, and JR. The investigation was undertaken by HY and PKC, with methodology development by HY, PKC, and PC. VT and JR provided supervision throughout the project. Validation was performed by HY and PKC, and visualizations were created by HY, PKC, and PC. The original draft of the manuscript was written by HY, PKC, and PC, with review and editing contributions from HY, PKC, PC, HL, SS, VT, and JR.

## Supporting Informations

### A Equations

#### Response Index

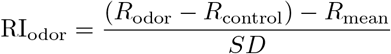

#### Cosine Similarity

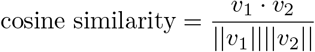

The · denotes the dot product, || · || denotes the norm of a vector.

### B Hyperparameters of STAN-Flow

Throughout the study we used a window size [Δ] = 20 msec, our data has a resolution of 1 msec, so 20 msec resulted in a dimension with size 20. The LSTM encoder has 2 layers of LSTM units, each has 4 hidden dimension. The output dimensions of the embedding layers for the attention module are 10. The stimuli are one-hot encoded. For the normalizing flow, we applied the RealNVP architecture [27] to construct a normalizing flow with 4 coupling layer blocks, each with a hidden dimension size of 64. The activation function for the s net is *tanh* and the activation for t net is *softplus*. To train STAN-Flow, we used an AdamW optimizer with a learning rate of 1e-5.

### C Additional Result

In this section of the supporting information, we include additional results including testing different parameters for pairwise synchronization methods, Response index matrix for other preparations; testing spatial attention’s performance across preparations; and exploring the PN, LN interaction with stimuli that has altered odor ratios.

#### C.1 Identification of the location of probes in the AL

Here, we use the approach to acquire the ensemble recordings from the Antennal lobe (AL) via microelectrode silicon probes and visualizing where location of these probes in the AL. We also quantify the number of units (neurons) obtained from these ensemble recordings.

### C.2 Testing different parameters for Ensemble Synchronization and Kernel Binless Methods

In this subsection, we test the Ensemble Synchronization index and kernel binless methods across multiple preparations with a wide range of hyperparameters. We ran similar 2D TSNE analyses as in section 4.2. Specifically in Fig S2, we compute ensemble synchronization over two different periods (0-500 msec, 0-1000 msec after onset of stimuli), and different binsize (5, 10, 20 msec). Limited clustering can be observed across different subjects and different parameters for the Ensemble synchronization index method.

For the Kernel binless method, we tested *τ* ∈ [2, 3, 4, 5, 6 msec] used to tune the exponential kernel. There is, however, no obvious separation between the behavioral and non-behavioral stimuli (Fig S3).

### C.3 RI of other preparations

We show the STAN-Flow generated and real Response index (RI) across different preparations in Fig S4. The response index matrices between generated and real spike trains are similar across 4 different preparations. Indicating that the STAN-Flow’s generative performance is consistent across preparations.

### C.4 Spatial attention performance on other preparations

We also conducted similar analyses in section 4.2 across different preparations as well. Although the spatial attention does not cluster for these preparations, we show that the behavioral and non-behavioral stimuli are still linearly separable (see Fig S5).

### C.5 Interaction between PNs and LNs with altered stimuli ratio

We show the clustering result of different neuron types for stimuli with altered composition ratios. Fig S6A and S6B shows the clustering of the different concentrations of BEA in the behavioral mixture. We also showcase the RI of this experiment.

**Figure S1.**
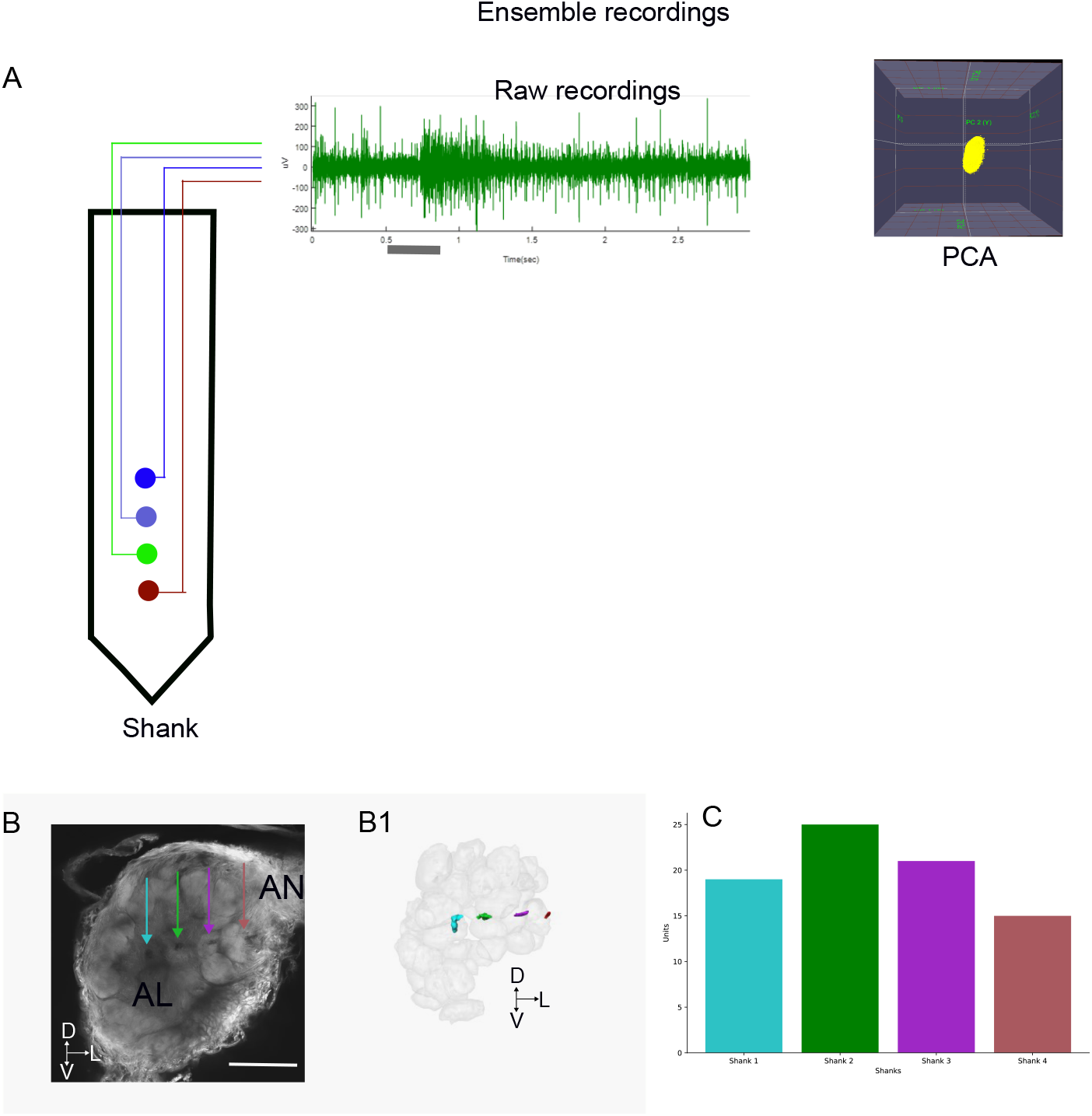
Schematic showing the acquisition of raw data from the antenna lobe (AL). (A) The location of the microelectrode silicon probes in the shank. Only one shank is used for the demonstration purpose. The green-colored silicon probe measures the odor-evoked response to the datura floral mixture. The gray bar represents the stimulation duration. The data is spike-sorted according to the PCA in the offline sorter. (B) The maximum intensity projection of the AL demonstrates the location of the probes in the AL. The 3D reconstruction of the probes and glomerulus that is impeded by these probes in the AL. Scale bar 100um. (C) The number of units obtained from the ensemble recording with four shanks. Note that the numbers are relatively higher in shanks 2 and 3. AN: Antennal nerve; D: Dorsal; V: Ventral; L: Lateral

**Figure S2.**
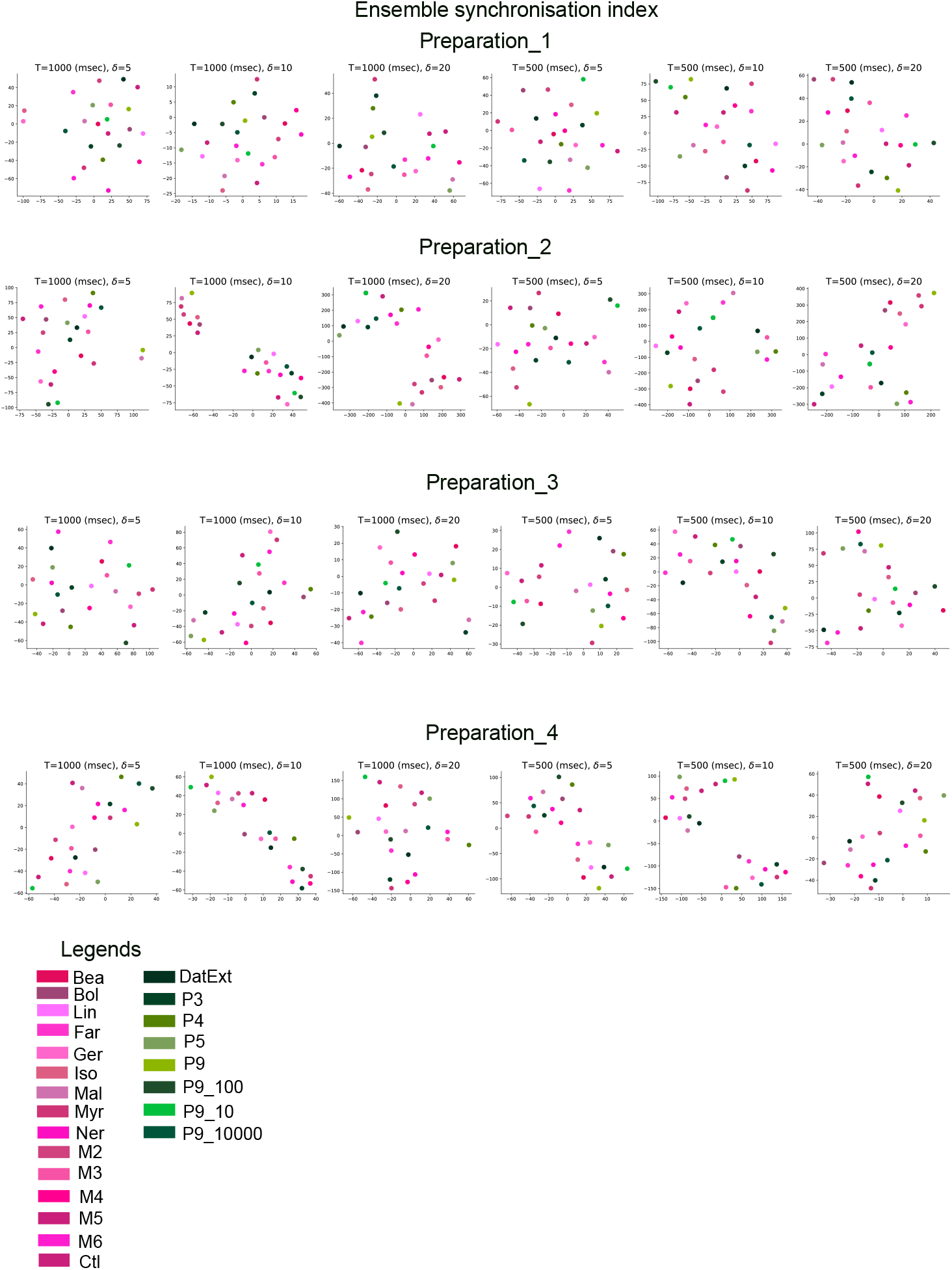
Two dimension TSNE analyses with the upper triangular matrices of the pairwise ensemble synchronization methods applied with different parameters on different preparations. *T* denotes the length of the stimulation segment that is analyzed, and δ denotes the binsize used in that specific computation of the ensemble synchronization index.

**Figure S3.**
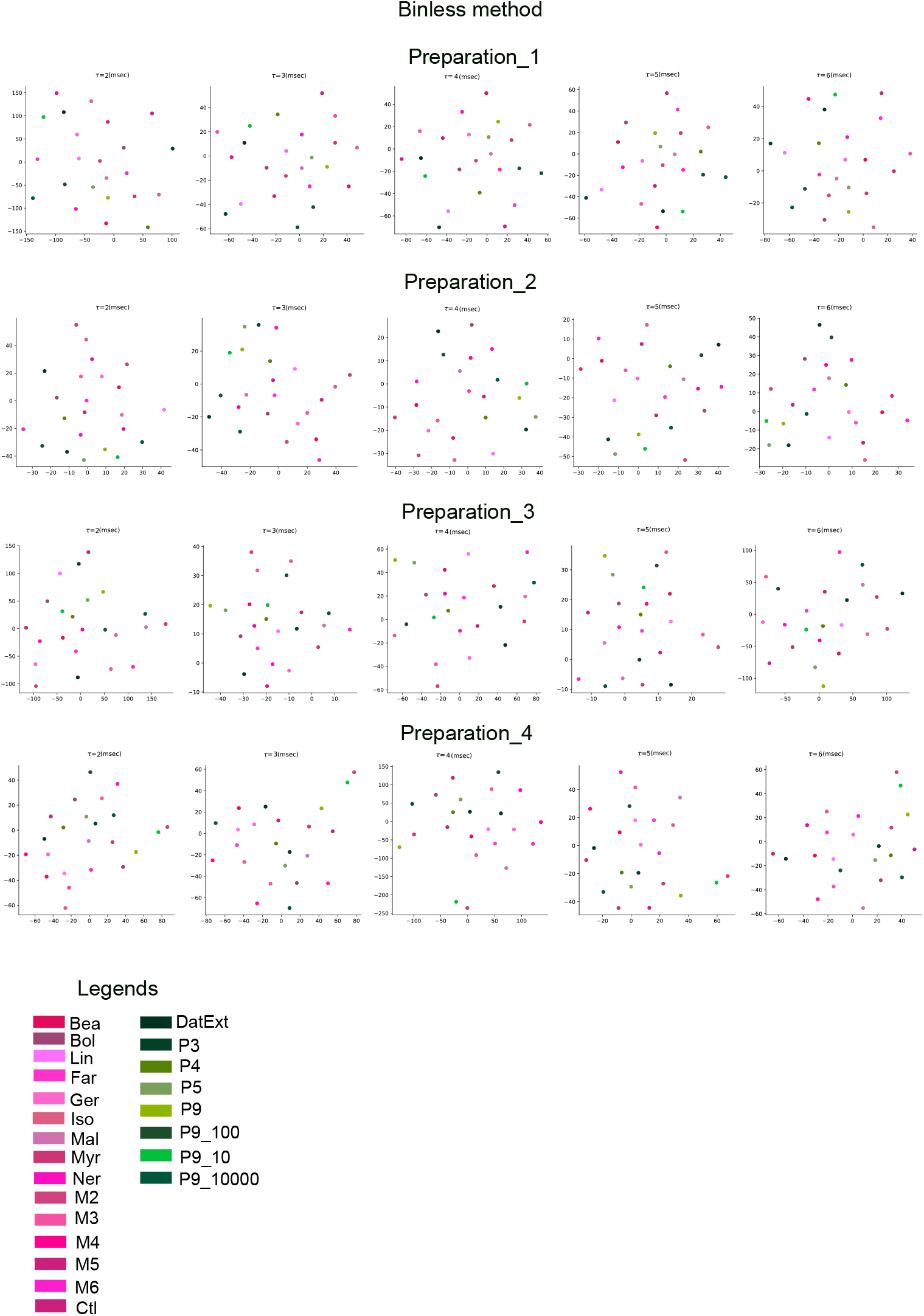
Two dimension TSNE analyses with the upper triangular matrices of the pairwise kernel binless methods applied with different parameters on different preparations. *τ* denotes the time constant parameter of the exponential kernel applied in the kernel binless method.

**Figure S4.**
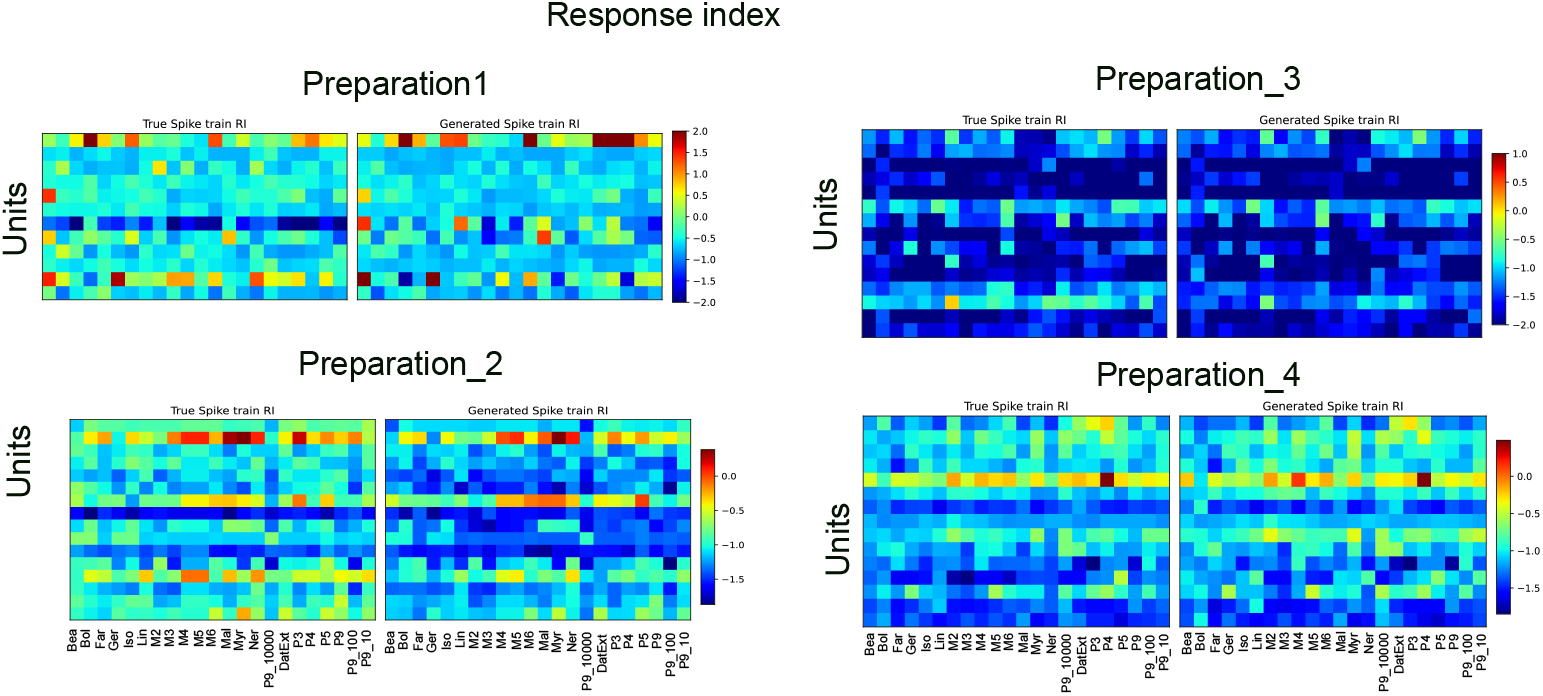
Response index of different preparations. Response index is calculated as shown in supporting information A.

**Figure S5.**
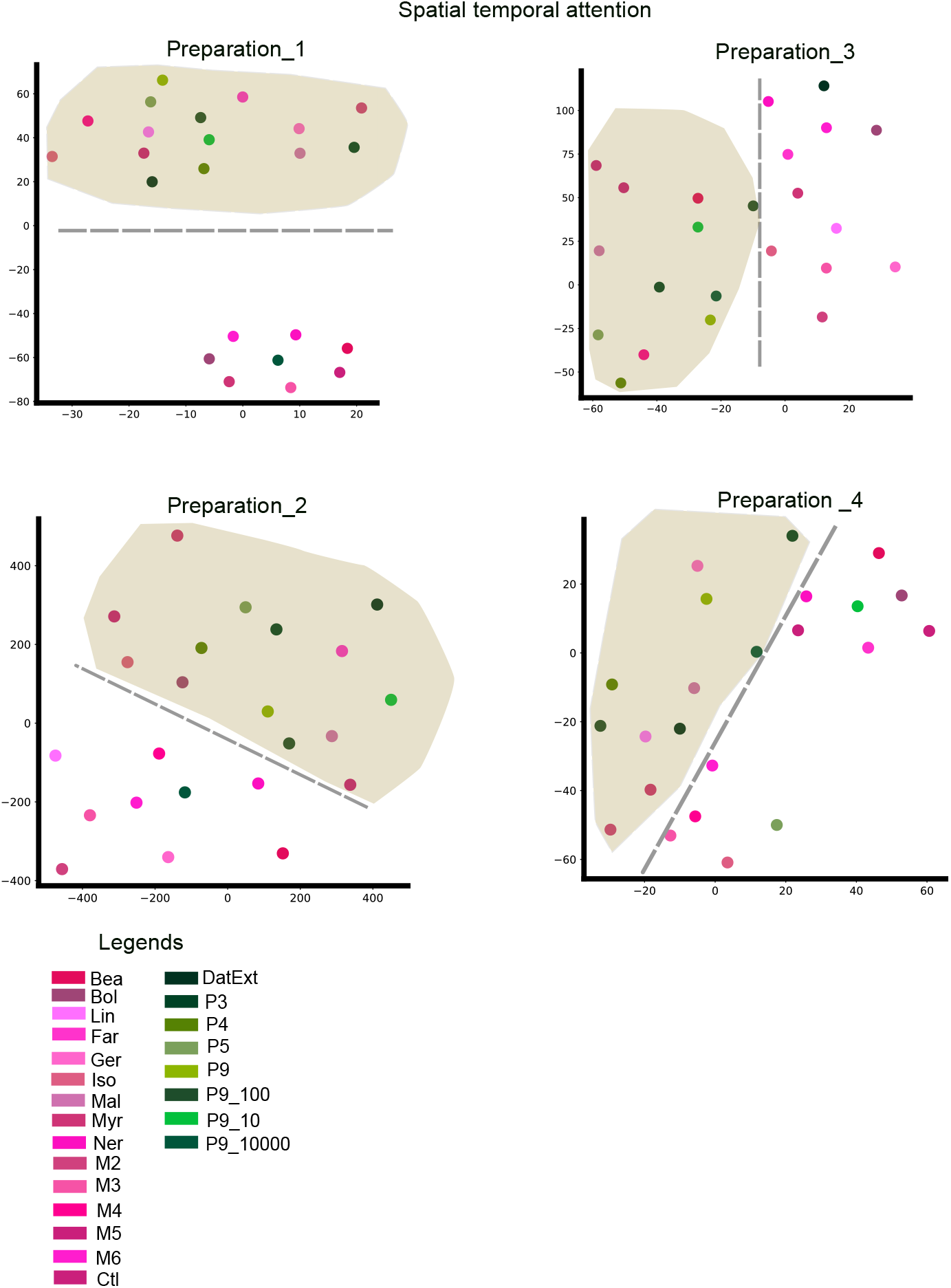
Spatial attention weights separate the behavioral stimuli (represented by different shades of green) from the non-behavioral stimuli (different shades of Magenta). With the gray line, we show that the different stimuli types remain linearly separable. The behavioral stimuli class is highlighted with a brown background.

**Figure S6.**
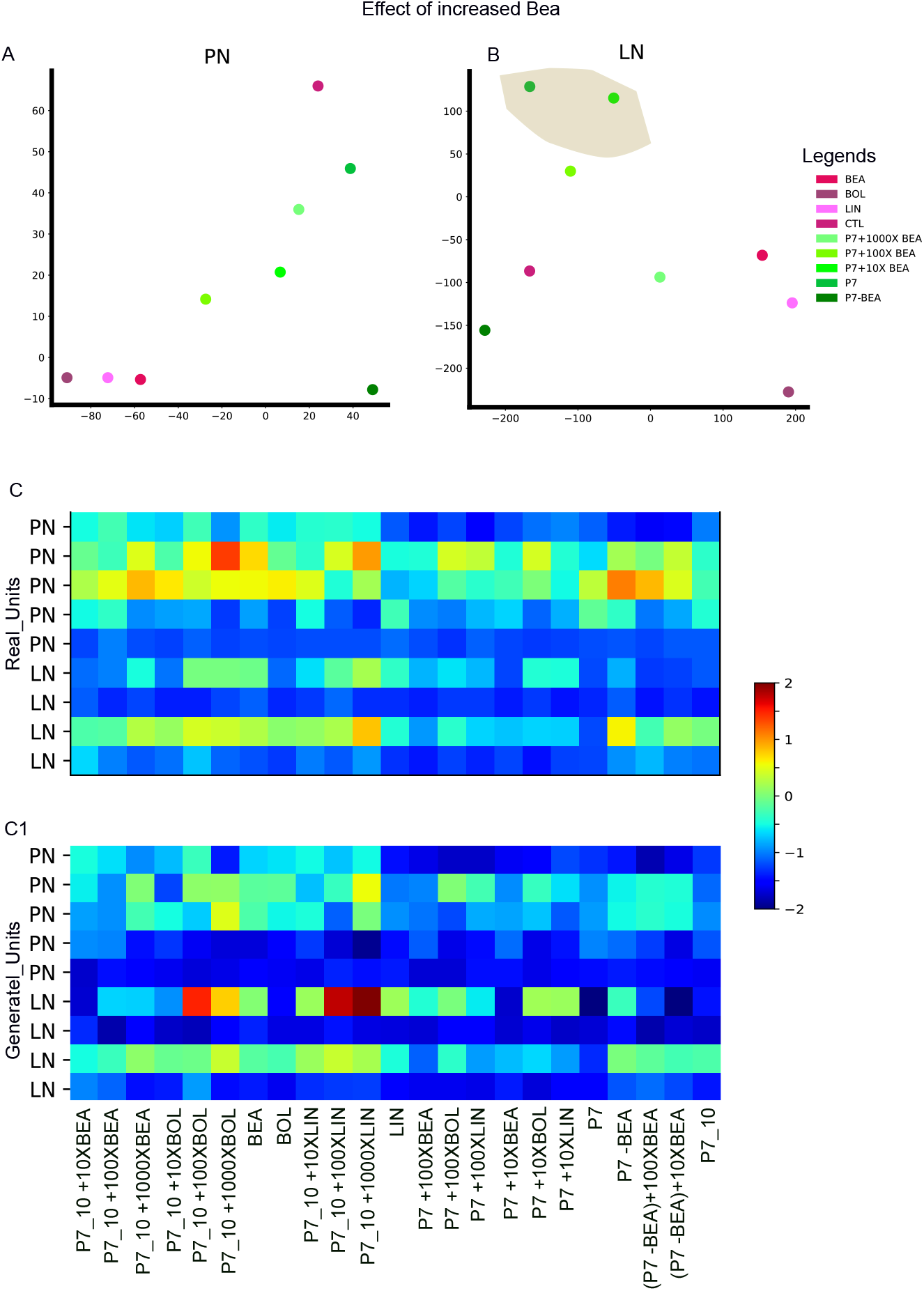
Clustering of the increased concentration of BEA with behavioral compound. The TSNE plot of the PNs (A) and LNs (B) demonstrates the clustering of the different concentrations of the BEA in the behavioral mixture (P7). The shaded brown area represents the clusterin of P7 mixture and P7 + 10 fold increase of BEA concentration (P7 + 10X BEA). (C, C1) Unit responses of 9 units to the different volatile stimuli tested and plotted as color-coded response matrices across all units (rows 1–9) and different volatile stimuli for the real units (C) and generated units (C1). Each units were classified into projection neurons (PNs) and Local interneurons (LNs). Note that the response index was affected when the behavioral odor (either BEA, BOL, or LIN) was increased in the mixture of the behavioral compounds diluted by 10-fold (P7 10). The effect was greater when Linalool concentration was increased by 1000 folds in the LN. Different component without BEA was also tested. The control (mineral oil) was used and is not shown in the plot.

## Footnotes

1 The KS-Test will reject the null hypothesis if two distributions are statistically different

2 Also known as the test of equivalence for two independent samples; the TOST will reject the null hypothesis if the means of the two distributions are within the *±*Δ region. We choose Δ = 2msec

